# Resistance to malaria through structural variation of red blood cell invasion receptors

**DOI:** 10.1101/083634

**Authors:** Ellen M. Leffler, Gavin Band, George B.J. Busby, Katja Kivinen, Quang Si Le, Geraldine M. Clarke, Kalifa A. Bojang, David J. Conway, Muminatou Jallow, Fatoumatta Sisay-Joof, Edith C. Bougouma, Valentina D. Mangano, David Modiano, Sodiomon B. Sirima, Eric Achidi, Tobias O. Apinjoh, Kevin Marsh, Carolyne M. Ndila, Norbert Peshu, Thomas N. Williams, Chris Drakeley, Alphaxard Manjurano, Hugh Reyburn, Eleanor Riley, David Kachala, Malcolm Molyneux, Vysaul Nyirongo, Terrie Taylor, Nicole Thornton, Louise Tilley, Shane Grimsley, Eleanor Drury, Jim Stalker, Victoria Cornelius, Christina Hubbart, Anna E. Jeffreys, Kate Rowlands, Kirk A. Rockett, Chris C.A. Spencer, Dominic P. Kwiatkowski, Malaria Genomic Epidemiology Network

## Abstract

*Plasmodium falciparum* invades human red blood cells by a series of interactions between host and parasite surface proteins. Here we analyse whole genome sequence data from worldwide human populations, including 765 new genomes from across sub-Saharan Africa, and identify a diverse array of large copy number variants affecting the host invasion receptor genes *GYPA* and *GYPB*. We find that a nearby reported association with severe malaria is explained by a complex structural variant that involves the loss of *GYPB* and gain of two hybrid genes, each with a GYPB extracellular domain and GYPA intracellular domain. This variant reduces the risk of severe malaria by 40% and has recently risen in frequency in parts of Kenya. We show that the structural variant encodes the Dantu blood group antigen, and therefore a serologically distinct red cell phenotype. These findings demonstrate that structural variation of red blood cell invasion receptors is associated with natural resistance to *P. falciparum* malaria.

## Main text

Malaria parasites cause human disease by invading and replicating inside red blood cells, which can lead to life-threatening complications that are a major cause of childhood mortality in Africa (1, 2). The invasion of red blood cells is orchestrated by the specific binding of parasite ligands to erythrocyte receptors (3), a stage at which genetic variation could influence the progression of infection. Indeed, a human genetic variant that prevents erythrocytic expression of the Duffy antigen receptor for chemokines (DARC), which is essential for invasion by *Plasmodium vivax*, is thought to have undergone a selective sweep in African populations, resulting in the present-day absence of *P. vivax* malaria across most of sub-Saharan Africa (4). In contrast the main cause of malaria in Africa, *P. falciparum*, has an expanded family of erythrocyte binding ligands targeting a different set of human receptors, most of which appear not to be individually required for invasion (5–7). Two of the earliest recognized invasion receptors are the glycophorins GYPA and GYPB, which are abundantly expressed on the erythrocyte surface and underlie the MNS blood group system (6, 8, 9). The high antigenic complexity of this system as well as rates of amino acid substitution and levels of diversity in African populations have led to speculation that this locus is under evolutionary selection due to malaria, but supporting epidemiological evidence is so far lacking (8, 10–12).

In a recent genome-wide association study (GWAS), we identified alleles associated with protection from severe malaria on chromosome 4, lying between *FREM3* and the cluster of genes encoding GYPE, GYPB and GYPA (13). Although the association signal did not extend to these genes and a functional variant was not identified, interpretation and further analysis of the association signal is inhibited by several factors. First, the GWAS samples were collected at multiple locations in sub-Saharan Africa, where levels of human genetic diversity are higher than in other parts of the world, and African populations remain relatively underrepresented in genome variation reference panels. Second, the glycophorin genes are in a region of segmental duplication that is difficult to characterize due to high levels of paralogy. Notably, the region is known to harbour multiple forms of structural variation that contribute to the MNS blood group system but have not been characterized by next generation sequence data (14, 15). Here we aim to capture additional variation in sub-Saharan African populations, including structural variation, to determine the underlying architecture of the association signal in this region.

### An African-enriched reference panel in the glycophorin region

We began by constructing a reference panel with improved representation of sub-Saharan African populations from countries where malaria is endemic. We performed genome sequencing of 765 individuals from 10 ethnic groups in the Gambia, Burkina Faso, Cameroon and Tanzania, including 207 family trios (100 bp paired end (PE) reads, mean coverage 10x; (16); **Tables S1** and **S2**). We focused on a region surrounding the observed association signal (chr4:140Mb-150Mb; GRCh37 coordinates). Genotypes at single nucleotide polymorphism (SNPs) and short indels in the region were called and computationally phased using published methods (17–19) and combined with Phase 3 of the 1000 Genomes Project (20) to obtain a reference panel of 3,269 individuals, including 1,269 Africans and a further 157 individuals with African ancestry (**Fig. S1** and **Tables S1** and **S3;** (16)). We imputed variants from this panel into the published severe malaria GWAS dataset comprising 4,579 cases of severe malaria and 5,310 population controls from the Gambia, Kenya and Malawi and tested for association as described previously (13, 16). The signal of association, formerly identified and replicated at SNPs lying between *FREM3* and *GYPE*, extends over a region of at least 700 kb, and includes linked variants within *GYPA* and *GYPB*, where association is only apparent with the additional African reference data (**Fig. S2**).

### Identification of copy number variants

We next assessed copy number variation in the glycophorin region (defined here as the segmental duplication within which the three genes lie; chr4:144,706,830-145,069,066; (16)) for the sequenced reference panel individuals. The high level of sequence identity between the duplicate units presents a considerable challenge for short read sequence analysis due to ambiguous mapping (**Fig. 1b**) (21). We therefore focused on changes in read depth at sites of high mappability. Specifically, we developed a hidden Markov model (HMM) to infer the underlying copy number state for each individual in 1600 bp windows using read depth averaged over sites above a set mappability threshold, and excluded windows with too few (<400) such sites. We adjusted for systematic variation along the genome with a window-specific factor computed across individuals. We then grouped individuals carrying similar copy number paths to assign copy number variant (CNV) genotypes (16).

**Fig. 1.**
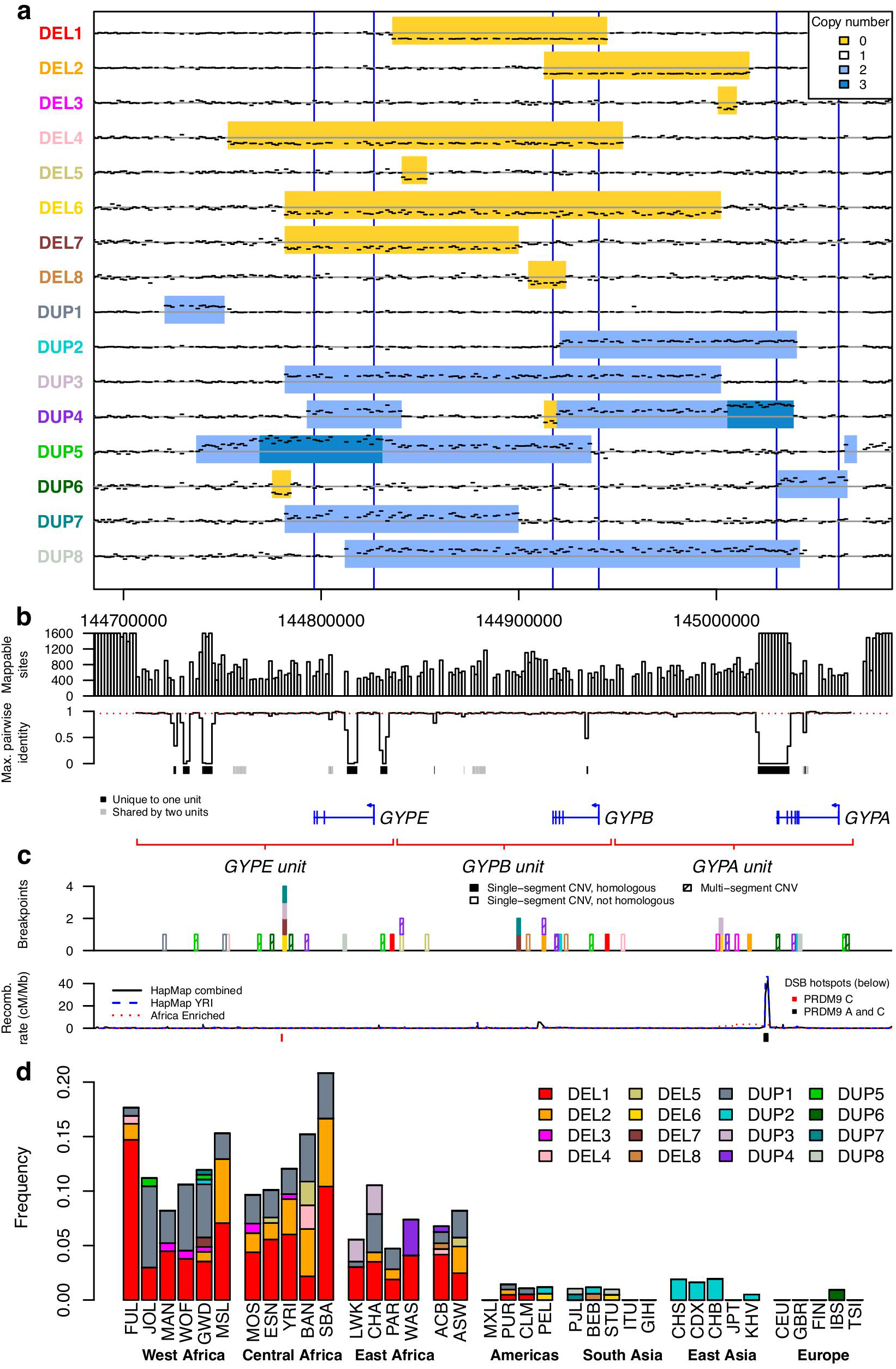
Copy number variants in the glycophorin region. **(a)** Sequence coverage in1600 bp windows and copy number for CNVs identified in ≥2 unrelated individuals. Black dashes show the mean sequence coverage across heterozygous carriers, normalized by the mean coverage of those individuals outside this region and the bin-specific factor for each window (16). Individuals carrying multiple variants were excluded from the computation and only bins with at least 25% mappable sites are displayed, as were input to the HMM. A horizontal gray line indicates the expected coverage without copy number variation. The inferred CNVs are indicated with deletion in yellow, duplication in light blue, and triplication in dark blue. Blue vertical lines mark the locations of the three genes. **(b)** Mappability in the glycophorin region. Rows show: (i) the number of mappable sites in the 1600 bp windows used for copy number inference (windows with less than 25% mappable sites not shown); (ii) the maximum identity with the homologous locations in the segmental duplication in the same 1600 bp windows, as inferred from a multiple sequence alignment, with mean of 0.96 indicated with a red dashed line; (iii) sequences of at least 100 bp that are unique to one (black) or two (grey) out of the three segmentally duplicated units; (iv) protein-coding genes; (v) location of the segmentally duplicated units. **(c)** Positions of breakpoints of variants shown in **(a)**, coloured as the variant names in **(a)** and shaded by whether the variant has a single pair of homologous breakpoints, a single pair of non-homologous breakpoints, or is a multi-segment CNV. The recombination rate from LD-based recombination maps (48, 49) and locations of DSB hotspots (23) are annotated below. **(d)** Frequency of each CNV in the sampled populations, estimated from unrelated individuals. Populations are grouped based on geographical proximity; population abbreviations can be found in **Tables S1** and **S3**.

Across the 3,269 samples, we identified eight deletions and eight duplications that were found in ≥2 unrelated individuals (referred to below as non-singleton CNVs), as well as at least 11 singleton variants (**Fig. 1a**, **Fig. S3** and **Supplementary Text**). For reference, we label these variants by copy number type (DEL for deletion, DUP for duplication), and number them in order of frequency. To validate the CNV calls we analysed transmission in family trios and observed segregation as expected with few exceptions (**Table S4** and **Supplementary Text**). We also compared the CNV calls with the 1000 Genomes Project structural variant analysis (22), and found highly consistent copy number inference (98.8% of individuals have the same copy number call) but improved resolution of overlapping variants in our analysis (**Fig. S4** and **Supplementary Text**). Validation of the breakpoint of the most common variant (DEL1) by Sanger sequencing further confirmed the accuracy of the calls made by our HMM (**Supplementary Text**).

The variants ranged in length from 3.2 kb (the minimum possible with our method) to >200 kb and included deletions and duplications of entire genes. Loss of *GYPB* was a common feature, with five different forms of *GYPB* deletion among the non-singleton CNVs (**Fig. 1a**). Hybrid gene structures were another common feature, with two non-singleton CNVs predicted to generate *GYPB-A* or *GYPE-A* hybrids (**Fig. S5**). Some variants are predicted to correspond to known MNS blood group antigens while others have not previously been reported (**Supplementary Text**). Of the non-singleton CNVs, half (8/16) had a single pair of breakpoints in homologous parts of the segmental duplication, consistent with formation via non-allelic homologous recombination (NAHR; **Fig. 1c**). Of these, four share a breakpoint position, which coincides with a double-strand break (DSB) hotspot active in a PRDM9 C allele carrier ((23); **Fig. 1c** and **Fig. S6**).

CNVs in the glycophorin region were observed more frequently in Africa than other parts of the world (**Fig. 1d**). In this dataset, the combined frequency of glycophorin CNVs in African populations was 11% compared to 1.1% in non-African populations, and most of the non-singleton CNVs (13/16) were found in individuals of African ancestry. Among fourteen different ethnic groups sampled in Africa, the estimated frequency ranged from 4.7-21% with the highest frequencies in west and central African populations.

### Association with severe malaria

We sought to incorporate CNVs into the phased reference panel with the aim of imputing into our GWAS dataset. Computational phasing of CNVs is challenging, as published methods do not model CNV mutational mechanisms or non-diploid copy number at smaller variants within CNVs. To work around this, we excluded SNPs and short indels within the glycophorin region and relied on the trio structure of sequenced individuals to resolve haplotype phase between CNVs and flanking SNPs (**Figs. S7 and S8**; (16, 18)). We evaluated how well CNVs can be imputed by comparing CNV genotype calls from the HMM with re-imputed genotype probabilities in the 1046 unrelated African individuals in our reference panel, in a cross-validation framework (16). This analysis predicts reasonable imputation performance for the three highest frequency CNVs, DEL1, DEL2, and DUP1, as well as for DUP4 and the rarer DEL7 (correlation between true and re-imputed genotypes > 0.8; **Fig. S9**). The combined reference panel, which may be of use for imputation in other studies, is available online.

We used this panel to impute CNVs into the severe malaria GWAS samples and tested for association as before (16). One of the imputed CNVs, DUP4, is associated with decreased risk of severe malaria (odds ratio, OR=0.59; 95% CI 0.48-0.71, *P* = 7.4x10^-8^ using an additive model with fixed-effect meta-analysis across populations; **Fig. 2a**). Across populations, evidence for association at DUP4 is among the strongest of any variant in our data. Moreover, conditioning on the imputed genotypes at DUP4 in the statistical association model removes signal at all other strongly associated variants including the previously reported markers of association (e.g., *P*_*conditional*_=0.54 at rs186873296; Fig. 2f, **Fig. S10**). DUP4 has an estimated heterozygous relative risk of 0.61 (95% CI 0.50-0.75) and its genetic effect appears to be consistent with an additive model, although the low frequency of homozygotes makes it difficult to distinguish the extent of dominance (homozygous relative risk 0.31; 95% CI 0.09-1.06; n=23 homozygotes). Analysis of different clinical forms of severe malaria showed that DUP4 reduced the risk of both cerebral malaria and severe malarial anaemia to a similar degree (**Table S5**). While we noted some evidence of additional associations in the region, including a possible protective effect of DEL2 (OR=0.61; 95%CI=0.41-0.92, *P*=0.02), these results are compatible with a primary signal of association that is well explained by an additive effect of DUP4.

**Fig. 2.**
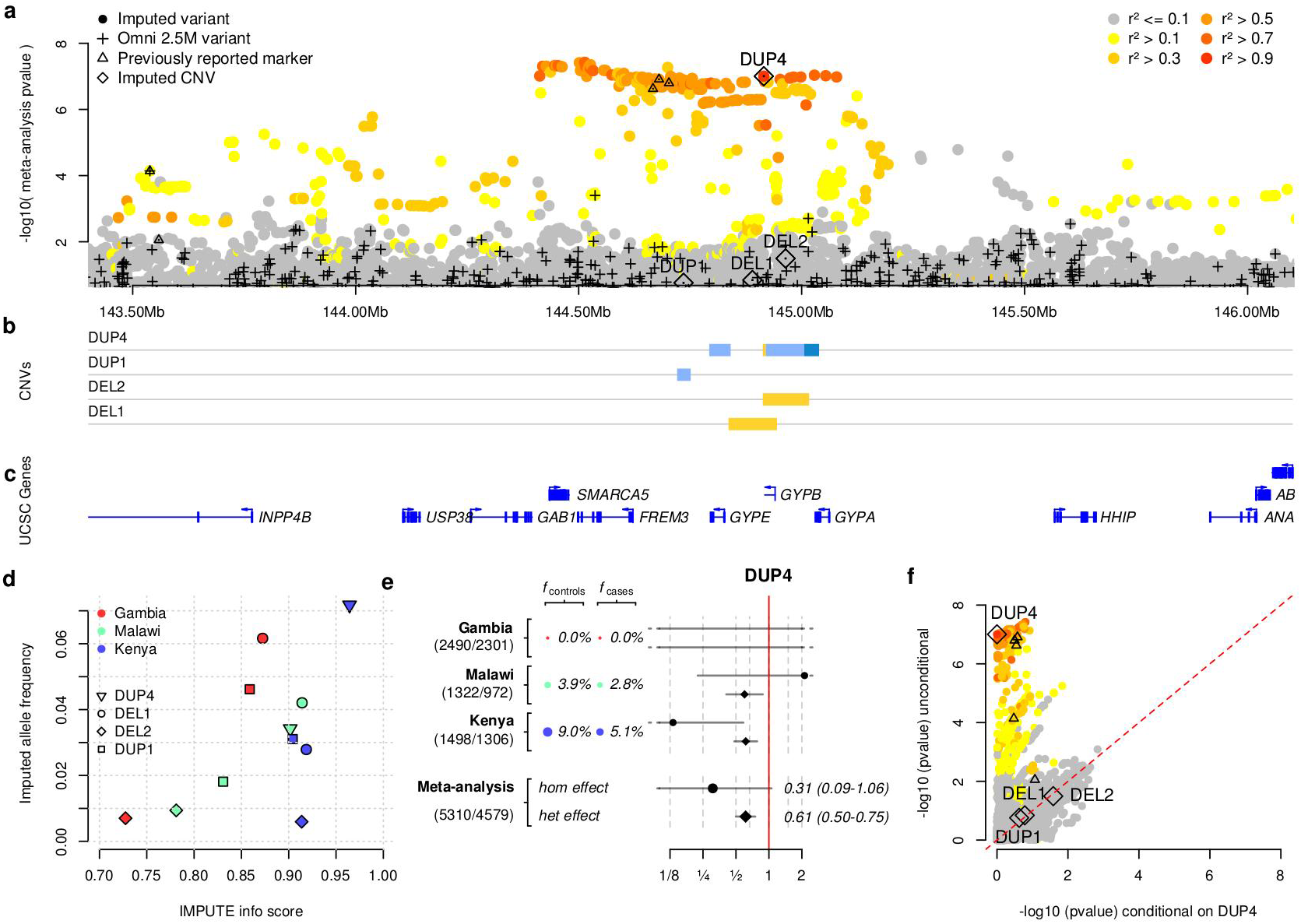
Evidence of association. **(a)** The evidence for association (-log10 *P*-value for association test, y axis) for SNPs, short indels, and CNVs across the glycophorin region (x axis). *P*-values are computed by meta-analysis across three African populations under an additive model of association. Points are coloured by LD with DUP4, as computed in east African reference panel populations. Directly typed SNPs are denoted with black plusses, and DEL1, DEL2, DUP1 and DUP4 are denoted with diamonds. Black triangles represent SNPs where the association signal was previously reported and replicated in further samples. **(b)** The copy number of DEL1, DEL2, DUP1 and DUP4, as in **Fig. 1a. (c)** Protein-coding genes in the region. **(d)** Comparison of expected imputed allele frequency (x axis) and IMPUTE info score (y axis), for the four annotated CNVs. **(e)** Detail of the evidence for association at DUP4. Coloured circles and text show the estimated allele frequency of DUP4 in population controls and severe malaria cases in each population. To the right is the odds ratio and 95% confidence interval for DUP4 heterozygotes (diamonds) and homozygotes (circles) relative to non-carriers. The bottom two rows represent effect sizes in a fixed-effect meta-analysis. Sample sizes (number of controls/number of cases) are denoted to the left. **(f)** Comparison of association test *P*-values conditioning on five principal components (y axis, as in panel **(a)**), and additionally conditioning on genotypes at DUP4 (x axis) for all variants in panel **(a)**.

DUP4 is imputed with high confidence in both east African populations (**Fig. 2d**), where it is at substantially higher frequency than in the reference panel (**Fig. S11**). To independently confirm the imputed DUP4 genotypes, we analysed SNP microarray data for intensity patterns indicative of copy number variation (**Fig. 3a**) using a Bayesian clustering model informed by the sequenced DUP4 carriers (**Supplementary Text**). Classification of GWAS samples was highly concordant with the imputed DUP4 genotypes in the east African populations (r^2^=0.97 in Kenya; r^2^=0.88 in Malawi). Surprisingly, both imputation and the microarray intensity analysis suggest there may be no copies of DUP4 present among the 4791 Gambian individuals in the GWAS. This large frequency difference places DUP4 as an outlier compared with imputed variants at a similar frequency in the Gambia or in Kenya genome-wide (empirical *P*=1.7x10^-3^ and *P*=5x10^*3^, respectively; **Fig. 4a**; (16)). Computation of haplotype homozygosity (**Fig. 4c**) provides evidence that DUP4 is carried on an extended haplotype (empirical *P*=0.012 for iHS (24, 25) compared with variants of similar frequency genome-wide; **Fig. 4d**) that may have risen to its current frequency in Kenya relatively recently. We note that DUP4 is also absent from all but two of the reference panel populations (**Fig. 1d**).

**Fig. 3.**
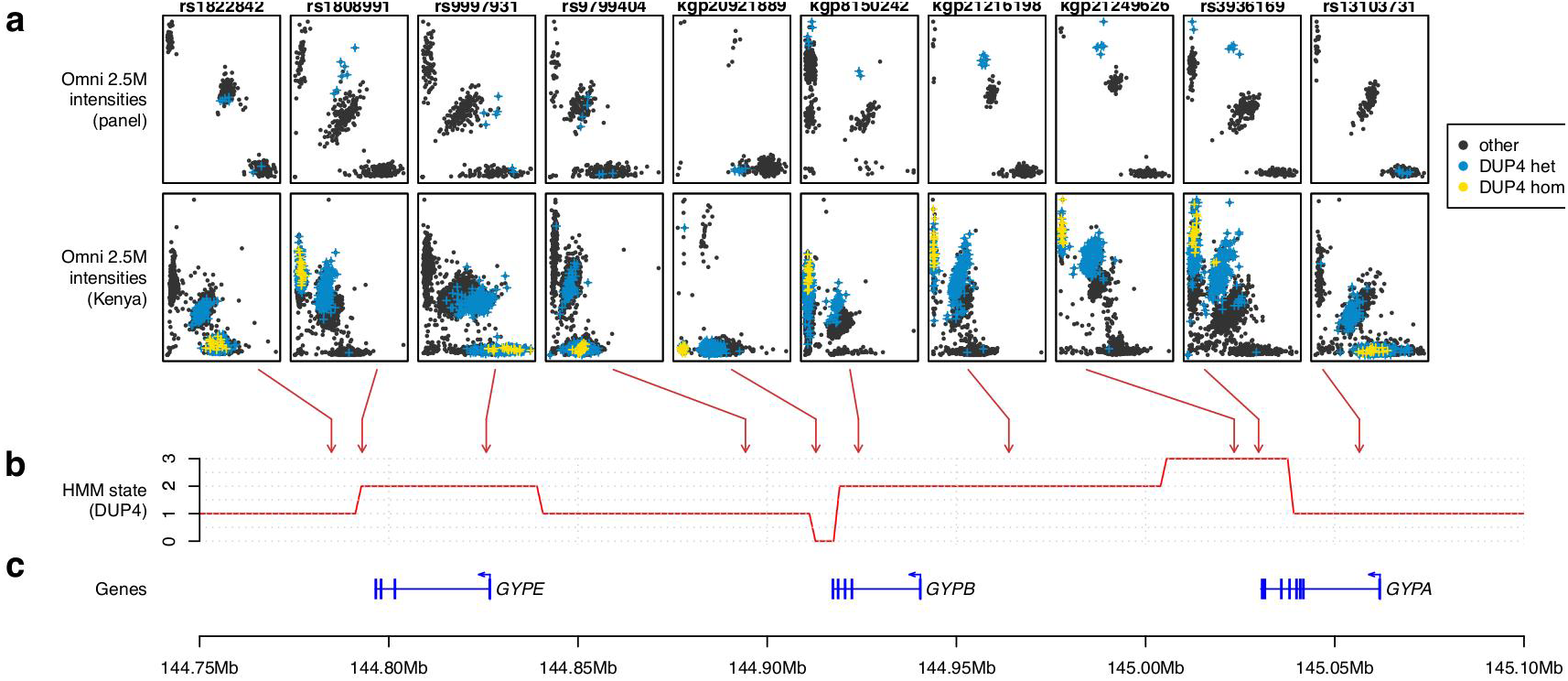
The effect of DUP4 on SNP array intensities. (**a**) Normalised Illumina Omni2.5M intensity values at selected SNP assays across the glycophorin region for reference panel individuals (top row; N=367 individuals from Burkina Faso, Cameroon, and Tanzania) and Kenyan GWAS individuals (second row, N = 3,142). Blue and yellow points represent individuals heterozygous or homozygous for DUP4 respectively, as determined by the HMM in reference panel individuals and by imputation in Kenya (genotypes with posterior probability at least 0.75). Arrows denote the mapping location of these SNPs. **(b)** HMM path for a single DUP4 individual. **(c)** Position of the glycophorin genes and exons.

**Fig. 4.**
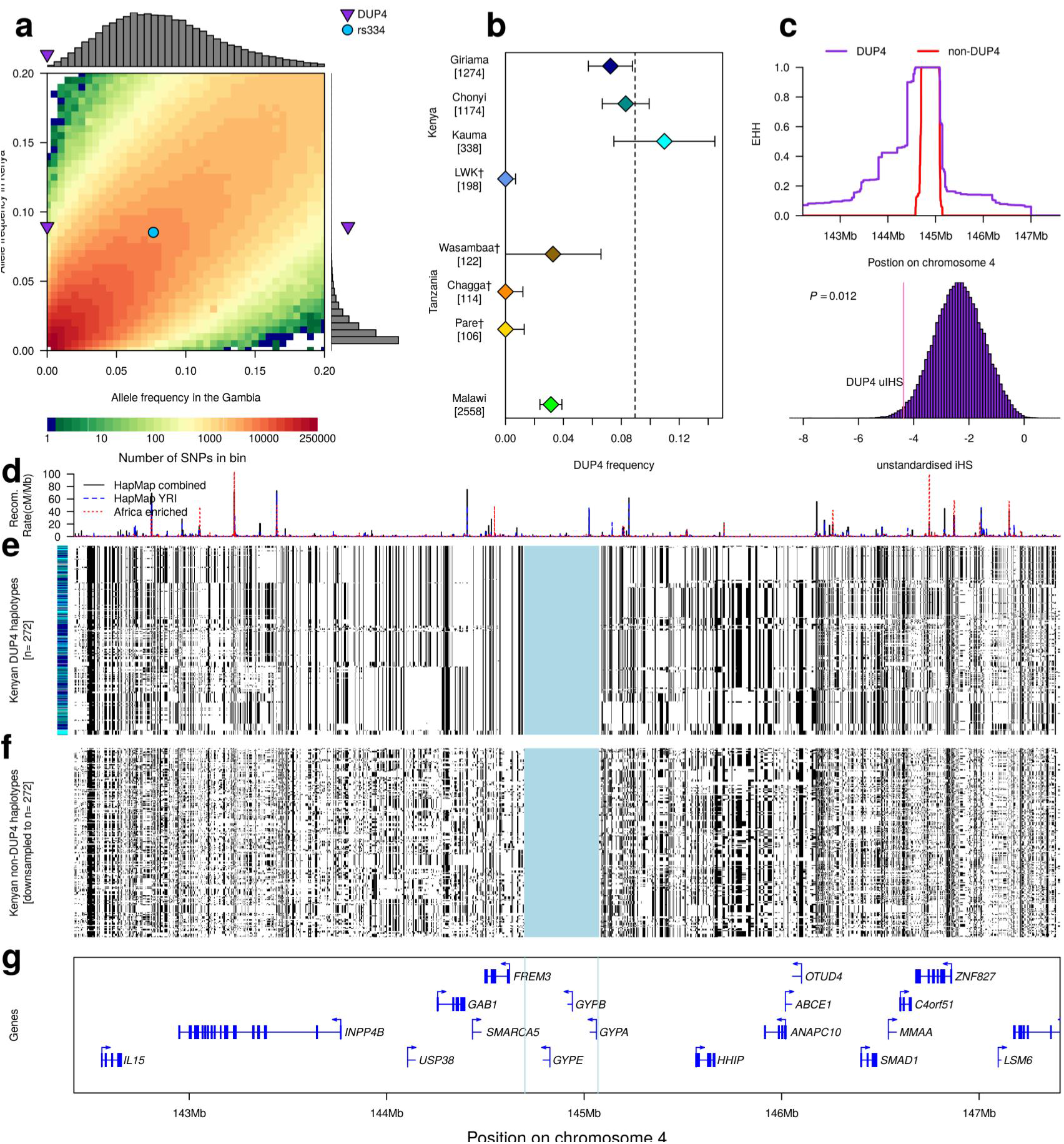
DUP4 frequency and haplotype homozygosity. **(a)** The empirical joint allele-frequency spectrum for population controls in the Gambia (x axis) and Kenya (y axis) in 0.5% frequency bins between 0 and 20% based on genome-wide imputed genotypes reported previously (13). Bins are coloured according to the number of variants, and the frequencies of DUP4 and rs334 are highlighted. The histograms show the marginal distribution of all SNPs in the Gambia in the same frequency bin as DUP4 in Kenya (8.5-9%, top) and the marginal distribution of all SNPs in Kenya in the same frequency bin as DUP4 in the Gambia (0-0.5%, right). **(b)** The estimated frequency of DUP4 in east African populations, shown as point estimates with 95% confidence intervals. The number of haplotypes sampled is given in brackets. Estimates are from population controls in the GWAS or, indicated by a dagger, from the HMM genotype calls in the reference panel. The overall frequency of DUP4 in the Kenyan controls (*f*_*Kenya*_ = 0.09) is shown as a dotted vertical line. **(c)** Extended haplotype homozygosity (EHH) computed outward from the glycophorin region for DUP4 haplotypes (purple) and non-DUP4 haplotypes (red) in Kenya, after excluding other variants within the glycophorin region. Below, the distribution of unstandardised iHS for all typed SNPs within 1% frequency of DUP4 in Kenyan controls. The position of DUP4 in this distribution is denoted by a red line, with empirical *P*-value annotated. **(d)** Recombination maps (48, 49) across the 142.5-147.5Mb region of chromosome 4. **(e)** The 272 haplotypes imputed to carry DUP4 in Kenya, clustered on 1 Mb extending in either direction from the glycophorin region, which is shaded in blue and was excluded from this analysis. The bar on the left depicts the population for each haplotype with colours as in panel **(b)**. **(f)** A random sample of 272 non-DUP4 haplotypes clustered on the same region. **(g)** Protein-coding genes.

### The physical structure of DUP4

The copy number profile of DUP4 is complex, with a total of six copy number changes that cannot have arisen by a single unequal crossover event from reference-like sequences (**Fig. 1a** and **Fig. 3b**). At the gene level, this copy number profile corresponds to duplication of *GYPE*, deletion of the 3’ end of *GYPB*, duplication of the 5’ end of *GYPB* and triplication of the 3’ end of *GYPA*. To begin to understand the functional consequences of DUP4, we sought to reconstruct the physical arrangement of this variant by pooling data across the nine carriers in the sequenced reference panel (eight Wasambaa individuals from Tanzania including three parent-child pairs, and a single African Caribbean individual from Barbados). First, analysis of coverage along a multiple sequence alignment of the segmental duplication corroborated the location of the six copy number changes from the HMM, with two pairs of breakpoints at homologous locations in the alignment (**Fig. S12**).

Next, we looked for sequenced read pairs spanning CNV breakpoints, which provide direct evidence of the structure of the underlying DNA. We identified read pairs that were mapped near breakpoints but with discordant positions (MQ>=1, absolute insert size > 1000 bp), including longer read data we generated for the 1000 Genomes individual who carries DUP4 (HG02554; 300 bp PE reads on Illumina MiSeq to 13x coverage; (16)). Discordant read pairs supported the connection between each pair of homologous breakpoints as well as between the remaining two breakpoints, which lie in non-homologous sequence (**Fig. 5a** and **Fig. S13**). Based on the combined evidence from copy number changes, discordant read pairs, and homology between inferred breakpoints, we generated a model of the DUP4 chromosome that contains five glycophorin genes (**Fig. 5b**).

**Fig. 5.**
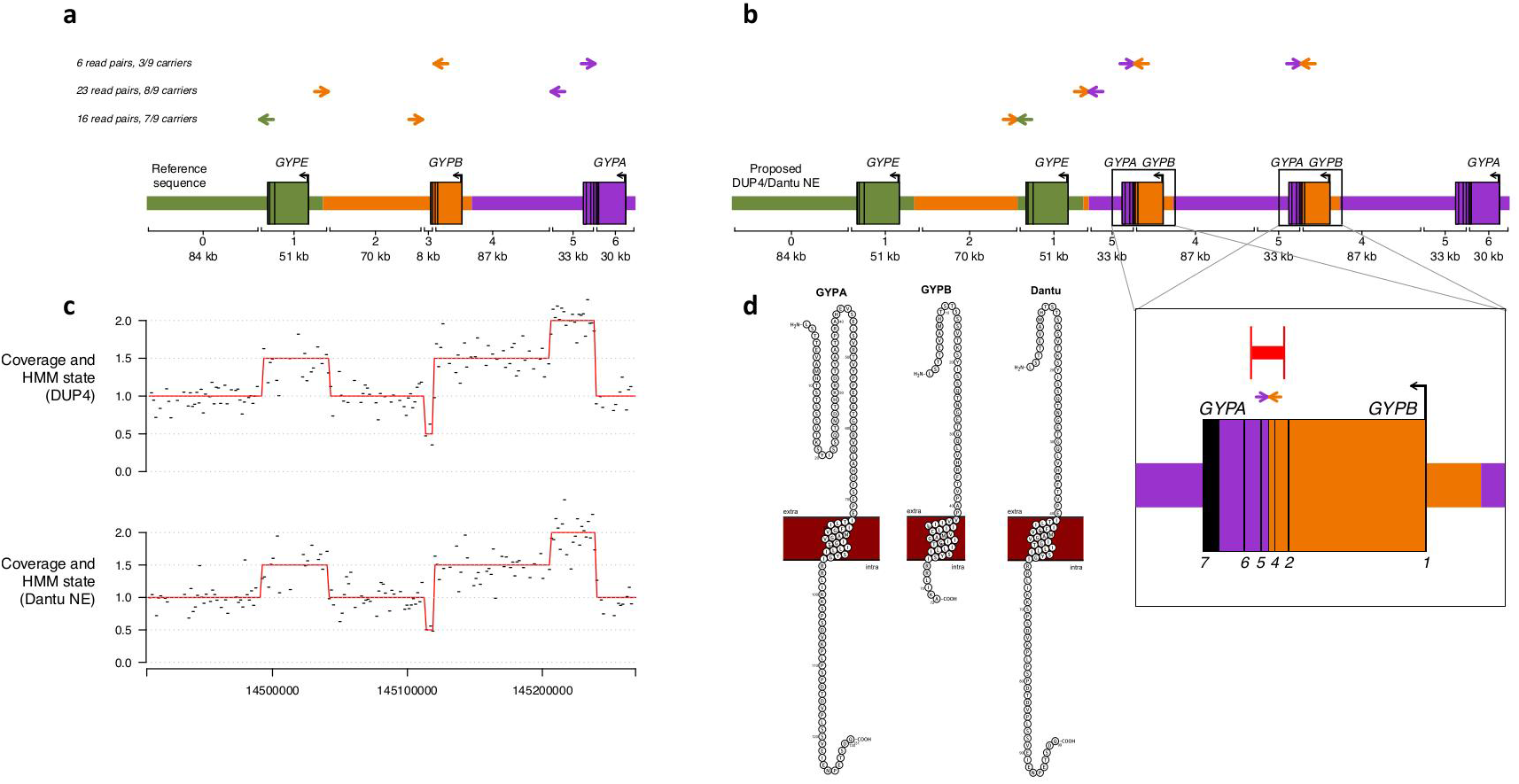
The structure of DUP4. **(a)** Discordant read pairs mapped near DUP4 copy number changes. Coloured arrows represent read pairs from heterozygous DUP4 carriers, with paired reads shown on the same horizontal line and the direction of the arrows depicting the strand and position as mapped to the human reference sequence. The number of such read pairs and distinct carriers in which they are found is given to the left. A schematic of the reference sequence is below with colors indicating the segmentally duplicated units. Brackets delineate the segments with different copy number in DUP4, which are numbered and labeled with their length to the nearest kb. **(b)** The structure of DUP4, inferred by connecting sequence at breakpoints based on sequence homology and discordant read pairs. Arrows depict the concordant positions of the read pairs in **(a)** on this structure, and the order of reference segments is shown below. Inset: detail of the inferred *GYPB-A* hybrid genes, indicating the positions of discordant read pairs (arrows), PCR primers (vertical red lines) and the resulting product (horizontal red line). **(c)** Normalized coverage in 1600 bp windows (black) and HMM path (red) for a DUP4 carrier (top) and for an individual serotyped as Dantu+ (NE type; bottom), on the same x axis as **(a)**. **(d)** Protein sequences of GYPA (NP_002090), GYPB (NP_002091), and the Dantu hybrid within the cell membrane depicting the extracellular, transmembrane, and intracellular domains as visualised with protter (50).

A prominent functional change on this structure is the presence of two *GYPB-A* hybrid genes, supported by several read pairs within intron 4 of *GYPA* and *GYPB* and the copy number profile. We confirmed the hybrid sequence by PCR-based Sanger sequencing of a 4.1 kb segment spanning the breakpoint (**Fig. 5b**, (16) and **Supplementary Text**). These data localize the breakpoint to a 184 bp section of *GYPA* and *GYPB* where the two genes have identical sequence (**Figs. S13** and **S14)**. If translated, the encoded protein would join the extracellular domain of GYPB to the transmembrane and intracellular domains of GYPA, creating a peptide sequence at their junction that is characteristic of the Dantu antigen in the MNS blood group system (**Fig. 5d**) (26, 27). Moreover, like DUP4, the most common Dantu variant (termed NE type, here referred to as Dantu NE) is reported to have two such hybrid genes and lack a full *GYPB* gene (28). We sequenced genomic DNA from an individual serologically determined to be Dantu positive, and of NE type (150 bp PE reads on Illumina HiSeq to 18x coverage; (16)) and analysed it using our HMM. The coverage profile and HMM-inferred copy number path, indistinguishable from those of DUP4 carriers, confirm identification of DUP4 as the molecular basis of Dantu NE (**Fig. 5c**).

To our knowledge, the full structure of this Dantu variant has not been previously reported (15). In addition to duplicate *GYPB-A* hybrid genes, these data reveal a duplicated copy of *GYPE* and the precise location of six breakpoints. Either complex mutational events or a series of at least four unequal crossover events are needed to account for the formation of this variant (confirmed by simulation; **Fig. S15**; (16)). However, we find no potential intermediates and no obvious relationship between DUP4 and other structural variant haplotypes in the present dataset (**Fig. S7** and **Supplementary Text**). Further analysis of discordant read pairs identifies a number of shorter discrepancies relative to the reference sequence that are consistent with gene conversion events (**Fig. S16**) and could be functionally relevant (e.g., **Fig. S17**).

## Discussion

Here we use whole-genome sequence data in combination with novel statistical methods to identify at least 27 CNVs in the glycophorin region that segregate in global populations. In this study, 14% of sub-Saharan African individuals carry a variant that affects the genic copy number relative to the reference assembly. Our description of these variants from genome sequence data complements the existing literature on antigenic variation associated with the MNS blood group system and offers novel insights. We find, for example, that the frequency of *GYPB* deletion is broadly commensurate with previous surveys of the S−s−U− blood group phenotype linked to absence of the GYPB protein, but the *GYPB* deletions in our data differ from the reported molecular variant (**Supplementary Text**; (15, 29–31)).

Of the array of glycophorin CNVs identified, one (DUP4) is associated with resistance to severe malaria and fully explains the previously reported signal of association (13). While there may be other functional mutations on this haplotype, we propose that the direct consequences of this rearrangement are likely to drive the underlying causal mechanism for resistance to severe malaria. DUP4 was not present in the 1000 Genomes Phase 1 reference panel (used in (13)), and exists as a singleton in the 1000 Genomes Phase 3 reference panel. Thus, as previously observed at the sickle cell locus (32), mapping of the association signal by imputation was only possible with the inclusion of additional individuals in the reference panel.

Through additional sequencing, we have shown that DUP4 corresponds to the variant encoding the Dantu+ (NE type) blood group phenotype, thus linking the predicted hybrid genes to a serologically distinct hybrid protein that is expressed on the red blood cell (27, 33). The few existing studies of Dantu+ (NE type) erythrocytes indicate high levels of the hybrid GYPB-A protein and lower levels of GYPA than wild type cells (33, 34) and a single study reports parasite growth to be impaired in vitro (35). Dantu NE is one of the many glycophorin variants that have been hypothesized to influence malaria susceptibility (11, 35–37), but the results here are the first evidence of a protective effect in natural populations.

These findings then raise the question of how DUP4 protects against malaria. GYPA and GYPB are exclusively expressed on the erythrocyte surface and are targeted by parasites during invasion (6, 7). *P. falciparum* EBA175 binds to the extracellular portion of GYPA (38), which is preserved in DUP4. *P. falciparum* EBL1 binds to the extracellular portion of GYPB (37) which is duplicated in DUP4 but joined to intracellular GYPA. The significance of the extra copy of *GYPE* or the absence of full *GYPB* in DUP4 is uncertain, since *GYPE* is not known to be expressed at the protein level (8, 39), and there is no evidence that absence of *GYPB* alone confers protection (**Fig. 2a**). GYPA and GYPB are known to form homodimers as well as heterodimers in the red cell membrane (31), so these copy number changes could have complex functional effects. There are physical interactions between GYPA and band 3 (encoded by *SLC4A1*) at the red cell surface (40) and parasite binding to GYPA appears to initiate a signal leading to increased membrane rigidity (41). Thus the GYPB-A hybrid proteins seen in DUP4 could potentially affect both receptor-ligand interactions and the physical properties of the red cell membrane.

Previous surveys of the Dantu blood group antigen have indicated that it is rare ((31, 42–44); **Table S6**). We find that DUP4 is absent or at very low frequency outside parts of East Africa, with a frequency difference and extended haplotype consistent with a recent rise in frequency in Kenya. In contrast, the malaria-protective variant causing sickle-cell anaemia (rs334 in *HBB*), which is thought to be under balancing selection, has a similar frequency in both the Gambia and Kenya (**Fig. 4a**). One possibility for why DUP4 is not more widespread, given its strong protective effect against malaria, is that it has arisen only very recently without time for gene flow to facilitate its dispersion. Alternatively, this frequency distribution could be consistent with balancing selection, for example if it protects only against certain strains of *P. falciparum* that are specific to East Africa. The glycophorin region is near a signal of long-term balancing selection, and measures of polymorphism in both the human glycophorins and *P. falciparum* EBA175 have been suggestive of diversifying selection (10–12, 45, 46). Although apparently not directly related to these signals, current selection on DUP4 may represent a snapshot of the long-term evolutionary processes acting at this locus. Mapping the allele frequency of DUP4 across additional populations could help clarify the nature of selection.

Recent GWAS have confirmed three other loci associated with severe malaria (*HBB*, *ABO*, *ATP2B4*), all of which are also related to red blood cell function (13, 47). However, the association with *GYPA* and *GYPB* stands out by directly involving variation in invasion receptors. These receptors have been found to be non-essential in experimental models (7, 9), yet this result indicates important functional roles in natural populations. Intriguingly, there is marked variation among *P. falciparum* strains in preference for different invasion pathways in vitro (7); field studies that account for parasite heterogeneity and tests for genetic interactions may therefore be important in determining how DUP4 affects parasite invasion. The discovery that a specific alteration of these invasion receptors confers substantial protection provides a foundation for experimental studies on the precise functional mechanism, and may lead us towards novel parasite vulnerabilities that can be utilized in future interventions against this deadly disease.

## Acknowledgements

We thank all the study participants and the members of the MalariaGEN consortium. A list of researchers involved at each study site can be found at https://www.malariagen.net/projects/consortial-project-1/malariagen-consortium-members.

The MalariaGEN Project is supported by the Wellcome Trust (WT077383/Z/05/Z) and the Bill & Melinda Gates Foundation through the Foundations of the National Institutes of Health (566) as part of the Grand Challenges in Global Health Initiative. The Resource Centre for Genomic Epidemiology of Malaria is supported by the Wellcome Trust (090770/Z/09/Z). This research was supported by the Medical Research Council (G0600718; G0600230; MR/M006212/1). Chris C.A. Spencer was supported by a Wellcome Trust Career Development Fellowship (097364/Z/11/Z). The Wellcome Trust also provides core awards to The Wellcome Trust Centre for Human Genetics (090532/Z/09/Z) and the Wellcome Trust Sanger Institute (098051).

Eric Achidi received partial funding from the European Community’s Seventh Framework Programme (FP7/2007-2013) under grant agreement N° 242095 – EVIMalaR and the Central African Network for Tuberculosis, HIV/AIDS and Malaria (CANTAM) funded by the European and Developing Countries Clinical Trials Partnership (EDCTP). Thomas N. Williams is funded by Senior Fellowships from the Wellcome Trust (076934/Z/05/Z and 091758/Z/10/Z) and through the European Community’s Seventh Framework Programme (FP7/2007-2013) under grant agreement N° 242095 – EVIMalaR. The KEMRI-Wellcome Trust Programme is funded through core support from the Wellcome Trust. Carolyne Ndila is supported through a strategic award to the KEMRI-Wellcome Trust Programme by the Wellcome Trust (084538). Tanzania/KCMC/JMP received funding from MRC grant number (G9901439). The Malawi-Liverpool-Wellcome Trust Clinical Research Programme (MLW) is a Major Overseas Programme of the Wellcome Trust. Malcolm Molyneux was funded by a Wellcome Trust Research Leave Fellowship. Vysaul Nyirongo was supported on the MLW core grant.

We thank the staff of the WTSI Sample Logistics, Genotyping, Sequencing and Informatics facilities and the WTCHG High-Throughput Genomics core for their contributions to sample handling and generation and processing of sequence data.

The multiple sequence alignment of the three glycophorin segmental duplication units is available at XXX and the combined reference panel is available at XXX. The additional sequence data generated for HG02554 and sequence data for the Dantu+ (NE type) individual are available under accession numbers XXX and XXX.

